# PHENOCAUZ: Decoding the Molecular Basis of Human Symptoms from Mendelian Disease Biology

**DOI:** 10.64898/2026.03.26.714535

**Authors:** Hongyi Zhou, Jeffrey Skolnick

## Abstract

Clinical symptoms are the observable manifestations through which human disease is recognized, yet a general framework connecting these phenotypes to their underlying molecular mechanisms is lacking. We hypothesized that individual symptoms emerge from perturbations of characteristic sets of proteins and pathways and that Mendelian disorders, in which causal genes are often known, provide natural human perturbation experiments from which these relationships can be learned. Here we present PHENOCAUZ, a proteome-scale framework that maps symptoms to molecular determinants using Mendelian disease biology and predicts candidate symptom drivers beyond known Mendelian proteins. Across 7,152 symptoms and 4,222 Mendelian proteins, PHENOCAUZ yields strong enrichment of held-out symptom-associated proteins, with an estimated precision of approximately 0.60 among the top 20 predictions for well-characterized symptoms, and more than doubles the number of candidate symptom-associated proteins. Independent comparisons with curated resources support both protein-level predictions and, more strongly, their pathway context. Efficacious drug targets and predicted mode-of-action proteins frequently differ from inferred symptom drivers at the individual-protein level but converge with them at the pathway level, revealing a mechanistic distinction between proteins that generate a phenotype and proteins through which that phenotype can be therapeutically modulated. We further show that the resulting molecular symptom map can be used to identify proteins associated with severe adverse effects, improve target-based prediction of serious drug reactions, and prioritize candidate therapeutics across cancer and non-cancer diseases. PHENOCAUZ thus provides a scalable framework linking clinical phenotypes to molecular mechanisms and actionable therapeutic pathways.

## Introduction

Clinical symptoms are the observable manifestations through which diseases are recognized and managed^1^. Adverse drug reactions and therapeutic responses are likewise expressed as changes in patient phenotype^2^. Despite this central role in medicine, the molecular determinants of most symptoms remain incompletely defined. A molecular map connecting specific phenotypes to the proteins and pathways that generate them could provide a mechanistic bridge from clinical presentation to disease biology, therapeutic intervention and drug safety.

Mendelian diseases offer a distinctive route to constructing such a map because their causal genes are frequently known^3^. We reasoned that a pathogenic perturbation that produces a particular symptom in a Mendelian disorder can serve as a natural human perturbation experiment: the affected protein and its molecular context provide information about mechanisms capable of generating that phenotype. Related perturbations of the same protein or pathway may contribute to the same symptom in a complex disease or after pharmacological intervention. Mendelian genotype-phenotype relationships can therefore provide a principled training set for inferring the molecular architecture of symptoms beyond rare genetic disease.

Previous studies have constructed symptom-disease networks^4,5^, investigated symptom clusters^6^ and predicted adverse drug reactions^7–10^. These approaches have established useful statistical relationships among diseases, phenotypes and drugs, but generally do not distinguish proteins with evidence of a causal relationship to a phenotype from proteins that are merely associated with the same disease or pathway context. This distinction is important for translating symptom biology into mechanistic hypotheses and therapeutic strategies.

To address these issues, we introduce PHENOCAUZ, a framework that learns molecular features of human symptoms from Mendelian disease biology and generalizes them across the human proteome. PHENOCAUZ integrates Mendelian phenotype-protein relationships with pathway membership, biological processes and protein disease-propensity features to identify molecular pathways associated with individual symptoms and to rank additional candidate symptom drivers. We evaluate these predictions against independent curated protein and pathway resources and then ask a complementary therapeutic question: do efficacious drug targets coincide with symptom drivers, or do they instead modulate the same molecular pathways? Finally, we examine how this phenotype-to-mechanism map can support target de-risking, prediction of serious adverse drug reactions and prioritization of candidate therapeutics.

## Results

An overview of PHENOCAUZ and how it works is presented in Figure 1. Additional details of the methodology are provided in Materials and Methods.

**Figure 1.**
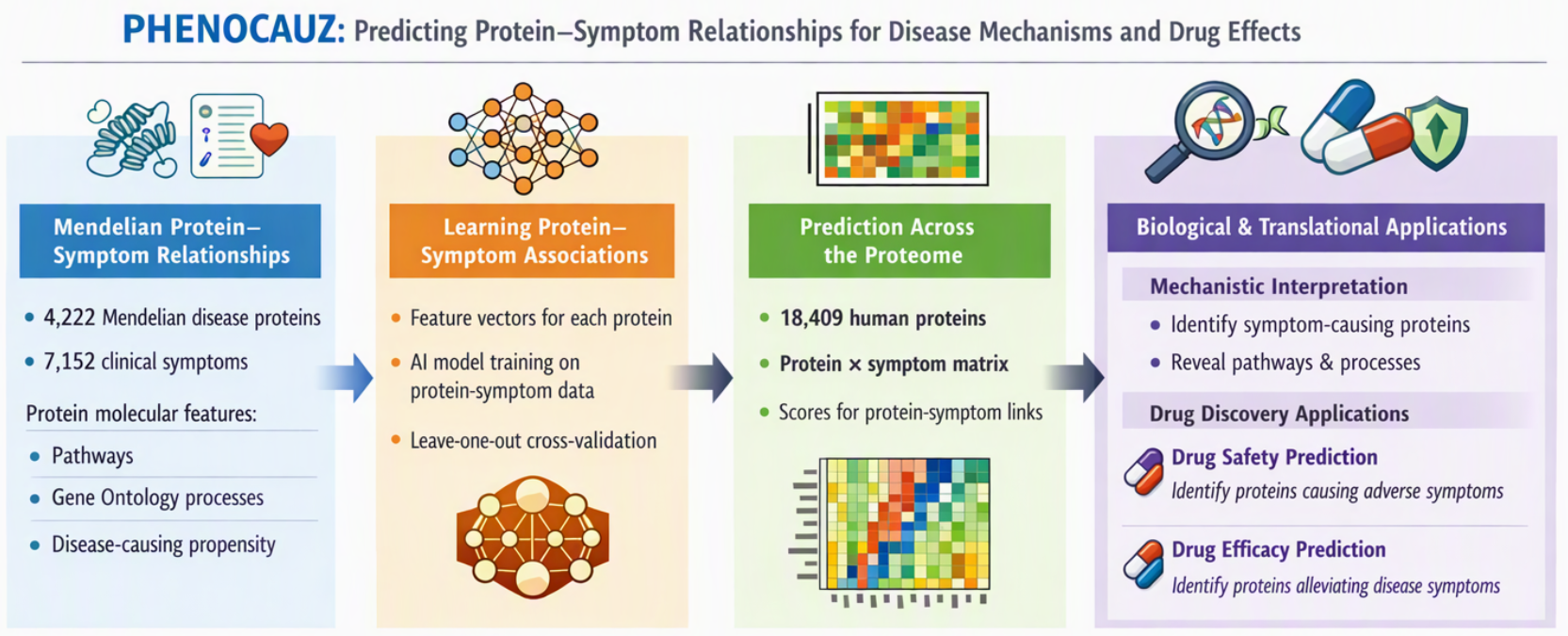
PHENOCAUZ framework for linking human symptoms to their molecular determinants. PHENOCAUZ leverages gene–phenotype relationships from Mendelian diseases to infer proteins and biological pathways responsible for human symptoms. Known Mendelian disease genes are first mapped to symptom–protein relationships, which are used to identify pathways and biological processes associated with specific clinical phenotypes. These molecular features are then used to train machine learning models that predict previously unrecognized symptom-causing proteins across the human proteome. The resulting symptom–protein map provides a molecular framework for interpreting disease phenotypes and enables applications in drug safety prediction and therapeutic target discovery.

### Construction of a symptom–protein map from Mendelian diseases

We first constructed a symptom–protein map by integrating Mendelian phenotype– gene relationship annotations^3^ with phenotype-symptom annotations obtained by PubMed text mining^11^. Mapping OMIM phenotype information to curated symptom taxonomies^2,5^ yielded 7,152 symptoms associated with 4,222 Mendelian proteins. These relationships form the initial training set for identifying molecular features associated with specific symptoms (the full lists of symptoms and protein-symptom causative relationships are in Supplementary Materials, Tables S1 and S2). The Developmental delay symptom has the most associated proteins (2,537), followed by Atrophy, with 2,261 proteins. The mean number of proteins causing a symptom is 153.2. The protein associated with the largest number of symptoms is TNF (1,973), followed by ITGB3 (1,966), while the mean number of symptoms associated with a protein is 259.5. The full list of proteins and their symptom frequencies are found in Supplementary Materials, Table S3.

### Molecular pathways associated with individual symptoms

Next, statistical analysis identified several Reactome pathways^12^ that were significantly enriched among the symptoms. Prominent among these are signal transduction, immune system pathways, metabolism, and developmental biology. These findings indicate that many symptoms arise from perturbations in identifiable molecular pathways. The top Reactome pathways are listed in Table 1 (the full list of symptom frequencies involving 1,238 pathways is in Supplementary Materials, Table S4, with detailed individual symptom’s top 20 lists in Supplementary Materials, Table S5). The top ranked pathway is Signal Transduction which is associated with 444 pain related symptoms, including abdominal, back and chronic cancer pain. Pain involves signal transduction that begins in the peripheral nervous system which is then interpreted by the central nervous system^13,14^. Depression, with 105 proteins involved in Signal Transduction, is supported by earlier studies^15^. “Weight increased” has 60 proteins involving the Signal Transduction pathway, which regulates energy homeostasis, appetite, glucose and fat metabolism, and adipose tissue development^16^. The second top ranked pathway is Immune System which is associated with 102 infection-related symptoms, including bacterial and Herpes virus infections. Examples of noninfectious symptoms in this pathway include Arthritis, Asthma, and Autoimmune diseases. Dementia has a top frequency of 161 proteins involving the Immune System, a result consistent with recent studies^17^. Other top pathways are Metabolism with 5,451 symptoms and Developmental Biology with 4,909 associated symptoms.

**Table 1.** Top 20 most frequent Reactome pathways causing symptoms.

| Number of symptoms within the top 20 pathways | ID | Name |
|---|---|---|
| 6240 | REACT:R-HSA-162582 | Signal Transduction |
| 6092 | REACT:R-HSA-168256 | Immune System |
| 5451 | REACT:R-HSA-1430728 | Metabolism |
| 5278 | REACT:R-HSA-168249 | Innate Immune System |
| 5006 | REACT:R-HSA-392499 | Metabolism of proteins |
| 4909 | REACT:R-HSA-1266738 | Developmental Biology |
| 4316 | REACT:R-HSA-372790 | Signaling by GPCR |
| 4264 | REACT:R-HSA-1280215 | Cytokine Signaling in Immune system |
| 3845 | REACT:R-HSA-1643685 | Disease |
| 3736 | KEGG:hsa01100 | Metabolic pathways |
| 3720 | REACT:R-HSA-74160 | Gene Expression |
| 3507 | REACT:R-HSA-109582 | Hemostasis |
| 3452 | REACT:R-HSA-422475 | Axon guidance |
| 3127 | REACT:R-HSA-449147 | Signaling by Interleukins |
| 3020 | REACT:R-HSA-382551 | Transmembrane transport of small molecules |
| 2983 | REACT:R-HSA-1280218 | Adaptive Immune System |
| 2899 | KEGG:hsa05200 | Pathways in cancer |
| 2054 | REACT:R-HSA-597592 | Post-translational protein modification |
| 2010 | REACT:R-HSA-556833 | Metabolism of lipids and lipoproteins |
| 1828 | KEGG:hsa04151 | PI3K-Akt signaling pathway |

### Prediction of symptoms caused by Mendelian proteins

We next asked whether molecular features learned from Mendelian phenotype-protein relationships could recover held-out symptom associations. To address this question, a Boosted Random Forest multilabel model was trained using pathway membership, biological processes and disease-propensity features. Leave-one-out cross-validation across 4,222 Mendelian proteins produced a mean enrichment factor of 8.5 across 7,152 symptoms; 82.7% of symptoms had enrichment factors greater than 1, and a mean per-symptom area under the Receiver Operating Characteristic (ROC) curve (AUC) of approximately 0.63. Because the intended use of PHENOCAUZ is to prioritize a short list of candidate proteins rather than to classify every protein-symptom pair, we also evaluated precision among the top-ranked predictions. For 77 symptoms with at least 1,000 known associated proteins, the mean top 20 precision was 0.52; for 23 symptoms with at least 1,500 known proteins, it increased to 0.60. These results indicate that the model substantially enriches relevant proteins near the top of the ranked list, particularly for well-characterized symptoms. Thus, it provides a prioritized list for subsequent experimental investigation.

### Prediction of symptom associations of proteins in complex diseases and Mendelian diseases without OMIM symptoms

We then generalized the model beyond known Mendelian symptom annotations. The proteome-wide analysis comprised 18,409 human proteins mapped to Reactome pathways^12^ and Gene Ontology biological processes^18^. Of these, 4,222 are Mendelian disease proteins with symptom relationships inferred from OMIM^3^ phenotypes; the remaining 14,187 proteins had no symptom annotations available to the algorithm. For a Mendelian protein, predictions were generated in leave-one-out fashion using the other 4,221 Mendelian proteins for training. For proteins without known Mendelian symptom annotations, the model was trained on all 4,222 annotated Mendelian proteins. We refer to proteins inferred in this second stage as PHENOCAUZ-predicted candidate symptom drivers, reserving causal terminology for relationships supported by Mendelian genetic evidence.

All predictions are combined into an 18,409 (proteins) × 7,152 (symptoms) score matrix. For each symptom, we extract known OMIM-derived proteins and additional predicted proteins with normalized score (Eq. (2)) > 0.32. This empirical selection threshold is close to the mean normalized score of top-ranked proteins for the 5,407 symptoms having more than 20 known proteins. In the held-out evaluation, mean top-20 precision was 0.52 for 77 symptoms with at least 1,000 known proteins and 0.60 for 23 symptoms with at least 1,500 known proteins (Figure 2). Known and predicted proteins are listed in Supplementary Materials, Table S6. On average, PHENOCAUZ predicts 139.5 additional proteins per symptom, compared with 153.2 known proteins; 32.4 of the additional proteins are Mendelian proteins not previously annotated with that symptom. Related approaches include a phenotype-similarity method that reported 0.131 precision across 3,069 rare diseases^19^ and PERADIGM, which evaluated three rare diseases^20^. Because the tasks, datasets and evaluation designs differ, these results do not support a direct numerical ranking against PHENOCAUZ; yet the above precision>0.5 is encouraging.

**Figure 2.**
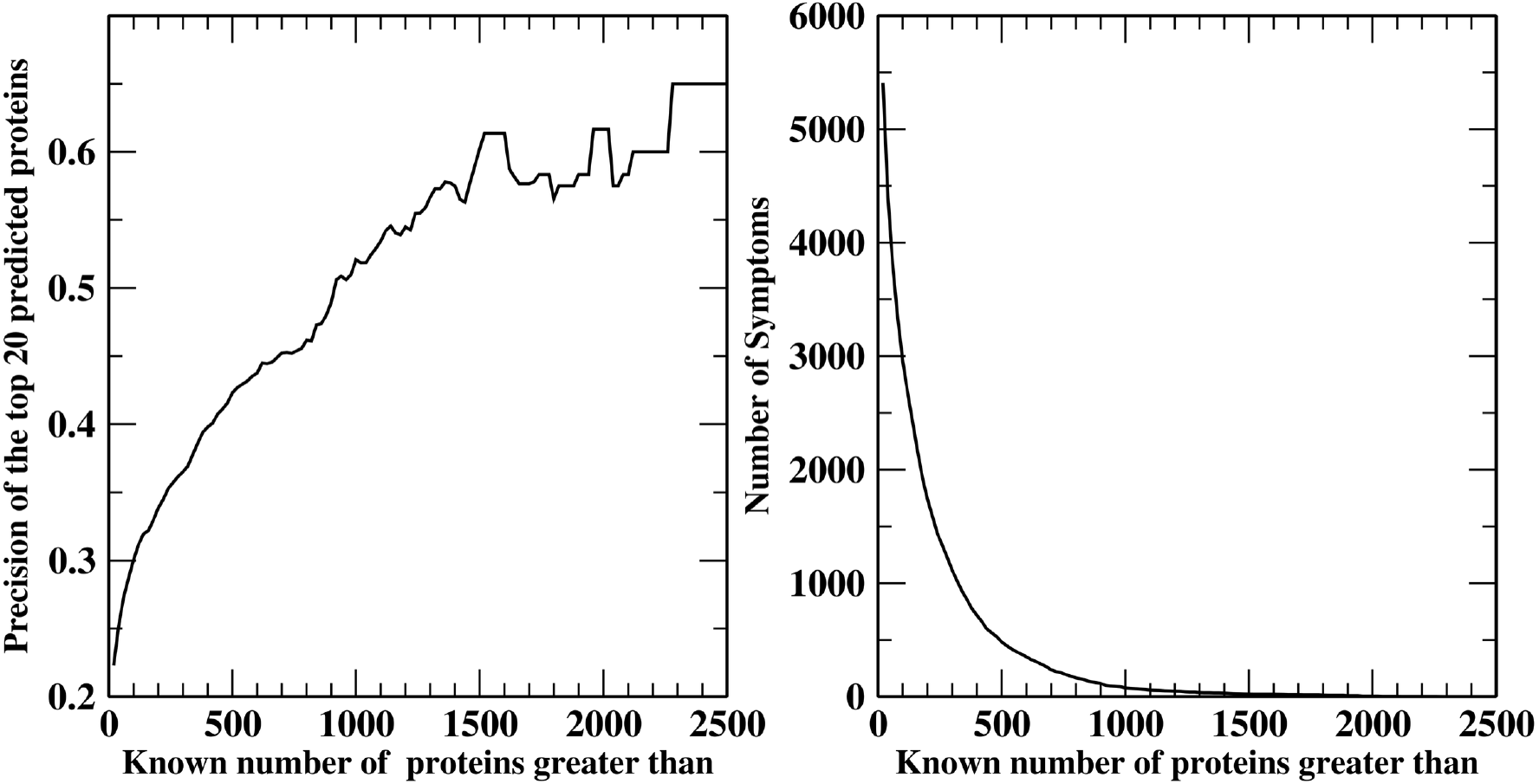
Mean precision among the top 20 predicted proteins versus the number of known proteins associated with a symptom (left), and the number of symptoms versus the number of known associated proteins (right). In the held-out evaluation, mean top 20 precision was 0.52 for symptoms with at least 1,000 known proteins and 0.60 for symptoms with at least 1,500 known proteins. These subset-specific estimates do not establish a universal upper or lower precision bound.

### Validation by literature curated symptom-protein and symptom-pathway associations

We evaluated PHENOCAUZ-predicted candidate symptom drivers against independent literature-curated resources using the Open Targets Platform^21^ and SympGAN^5^ which provide curated associations for independent comparisons. SympGAN contains symptom-protein associations that may include causal proteins as well as upstream or downstream pathway members, whereas the Open Targets Platform integrates broader disease-target evidence. Of the 7,152 PHENOCAUZ symptoms, 480 shared a UMLS_CUI identifier^22^ with SympGAN symptoms having curated protein associations, and 1,140 mapped to diseases represented in Open Targets Platform.

Direct protein overlap provided one level of independent support. Among 480 mapped symptoms, 352 (73%) showed significant overlap between Mendelian-derived proteins and SympGAN proteins (Fisher’s exact test, p < 0.05). Among 398 symptoms with at least one PHENOCAUZ-predicted candidate driver, 57 (14%) showed significant overlap with SympGAN. Against Open Targets Platform, significant overlap was observed for 1,013 of 1,140 symptoms (89%) for Mendelian-derived proteins and 566 of 971 symptoms (58%) for PHENOCAUZ-predicted proteins (Table 2). The lower direct overlap for predicted proteins is expected because these candidates are specifically intended to extend beyond currently annotated relationships; accordingly, direct overlap should be interpreted as external support rather than a complete estimate of prediction accuracy.

**Table 2.** Fractions of symptoms having significant protein/pathway overlaps (p-value < 0.05) between PHENOCAUZ and other datasets.

| **Protein overlaps** |  |  |
|---|---|---|
| Dataset | PHENOCAUZ derived proteins | PHENOCAUZ predicted proteins |
| SympGAN curated<sup>5</sup> | 352/480 (73%) | 57/398 (14%) |
| Open Targets Platform<sup>21</sup> | 1013/1140 (89%) | 566/971 (58%) |
| Efficacious drug targets<sup>7</sup> | 398/843 (47%) | 214/734 (29%) |
| Mode-of-action proteins<sup>26</sup> | 222/855 (26%) | 104/744 (14%) |
| **Pathway overlaps** |  |  |
| Dataset | PHENOCAUZ derived proteins | PHENOCAUZ predicted proteins |
| SympGAN curated | 449/471 (95%) | 361/394 (92%) |
| Open Targets Platform | 1122/1138 (99%) | 924/966 (96%) |
| Efficacious drug targets | 827/842 (98%) | 679/731 (93%) |
| Mode-of-action proteins | 825/839(98%) | 690/730 (95%) |

Because curated association proteins should nonetheless tend to participate in the same biological pathways as true causative proteins, we next evaluated pathway-level overlap between our known and predicted novel protein sets and other datasets’ proteins. The results for all symptoms with significant overlap are provided in Table 2. For derived protein pathways, 1,122/1,138 (99%) of symptoms have significant pathway overlaps (p-value < 0.05) with those of Open Targets Platform. For predicted proteins, the fraction of symptoms having significant pathway overlaps is 924/966 (96%). These results support the biological plausibility of the novel predictions even when direct protein literature support is limited.

### Efficacious drug targets and mode-of-action proteins are pathway-concordant with PHENOCAUZ symptom drivers

We next asked whether inferred symptom drivers, both known (OMIM-derived) and novel (PHENOCAUZ-predicted), are mechanistically consistent with efficacious therapies that alleviate the corresponding symptoms.

To enable drug-based validation, we mapped the symptom UMLS_CUIs (from SympGAN^5^ and SIDER4^2^) to Human Disease Ontology identifiers (DOIDs)^23^ used in our prior MEDICASCY framework^7^, which links indications to efficacious drugs based on their 2D chemical structure. Of the 7,152 symptoms, 855 mapped to 3,608 MEDICASCY indications (shared UMLS_CUIs). This then allowed retrieval of efficacious drug targets from DrugBank^24^ and other sources^25^ used in MEDICASCY training. Mode-of-action (MOA) proteins were derived from our previous work LeMeDISCO^26^ for the same 3,608 indications, which expanded the efficacious drug targets from databases to potential efficacious targets^26^.

Direct protein overlap between symptom drivers and efficacious drug targets was significant for only a subset of symptoms (47% for Mendelian-derived proteins and 29% for PHENOCAUZ-predicted proteins; see Table 2). In contrast, pathway concordance was pervasive: 827 of 842 symptoms (98%) showed significant overlap between pathways containing efficacious targets and those containing Mendelian-derived symptom proteins, and 679 of 731 (93%) did so for PHENOCAUZ-predicted candidate drivers (p < 0.05). Thus, proteins implicated in generating a phenotype and proteins targeted by effective therapies frequently differ at the individual-protein level while converging on the same molecular pathways. This separation between phenotype generation and therapeutic modulation is a central feature of the PHENOCAUZ framework.

As an illustrative case, obesity showed strong pathway concordance despite limited direct target overlap. The 720 known obesity-causative proteins span 1,664 unique Reactome pathways, whereas 238 obesity drug targets in the MEDICASCY set span 766 pathways^12^. These pathway sets share 668 pathways (p-value ≈ 0), including Signal Transduction and GPCR-related modules (Signaling by GPCR, GPCR downstream signaling, and G alpha (s) signaling events), which are among the most enriched pathways for obesity in our feature analysis. PHENOCAUZ also predicts 2,246 previously unannotated obesity proteins involving 2,037 pathways (the list of obesity drivers can be found in Supplementary Materials, Table S7). These share 734 pathways with obesity drug targets (p-value ≈ 0). Again, this highlights the same GPCR-related signaling architecture.

We then generalized this analysis to mode-of-action (MOA) proteins provided by LeMeDISCO^26^, which provides additional efficacious protein target sets using MEDICASCY-predicted drug indications^7^, FINDSITE^comb2.0^-based protein target inference^27^, and disease comorbidity-based validation^28^. As with efficacious drug targets, for known cases, direct protein overlap of the MOA proteins with PHENOCAUZ-derived symptom drivers was limited for 222/855 symptoms (26%). For novel predictions, significant protein overlap (p-value < 0.05) was observed for 104/744 (14%). In contrast, pathway-level overlap remained high. 825/839 (98%) pathways overlap for known proteins, and 690/730 (95%) pathways overlap for novel proteins with a p-value < 0.05 (Table 2). Across two independent therapeutic reference sets (efficacious drug targets^24,25^ and LeMeDISCO mode-of-action proteins^26^), PHENOCAUZ symptom drivers show strong agreement at the pathway level despite modest agreement at the individual protein level. Once again, these results support a model in which many effective therapies act through pathway-proximal modulators—upstream regulators, downstream effectors, or compensatory nodes—rather than directly targeting the root causal proteins. The obesity example further demonstrates that substantial therapeutic insight can be recovered from pathway concordance even when direct target overlap is minimal. Consequently, PHENOCAUZ provides a mechanistic bridge between symptoms, candidate causal proteins, and therapeutically actionable pathway context, enabling interpretable hypothesis generation for drug efficacy and safety (see below).

### Proteins linked to serious adverse effects

As a practical application, we used PHENOCAUZ symptom drivers to identify proteins whose perturbation is associated with serious adverse effects that can derail drug development. Using curated adverse-effect terms assembled in our prior MEDICASCY work^7^, we analyzed 9 severe adverse-effect categories: Neoplasm malignant, Sudden death, Cardiac failure, Cardiac failure congestive, Sudden cardiac death, Cardiac death, Haemorrhagic stroke, Death and Cerebral haemorrhage. Here, we additionally included the following symptoms containing the keyword “death”: Fetal death, Sudden infant death syndrome, Death neonatal, Anxiety death, Death rattle, Sudden unexplained death in epilepsy, and Apparent death. For each severe adverse-effect, we extracted the union of known (OMIM-derived) and predicted candidate symptom drivers (see Supplementary Materials, Table S6). In total, PHENOCAUZ implicates 6,400 proteins responsible for severe adverse effects (see Supplementary Material, Table S8). The most frequent protein possibly contributing to adverse events is SCN5A, a voltage gated sodium channel, which is a putative cause of 13 of the 16 severe symptoms. For example, mutations in SCN5A can cause Sudden cardiac death^29^.Another frequence protein is protein F5 (associated with coagulation) that causes 12 severe symptoms. Loss of cardiomyocyte-specific F5 leads to increased SCN1B expression (another voltage gated sodium channel) with enhanced susceptibility to stress, arrhythmias and sudden death^30^. Other representative examples include tumor suppressors CDKN2A and CYLD for malignant neoplasms^31,32^, ABCC6 for cardiac failure^33^, BARD1 for breast cancer risk^34^, TTF2 for thyroid cancer^35^. These results provide an interpretable list of potential “no-go” targets for which antagonism/inhibition may carry unacceptable risk. The full list of proteins associated with serious side effects can be found in Supplementary Materials, Table S8.

### Inferring serious drug adverse effects from binding targets

Given a drug’s targets, PHENOCAUZ enables inference of its likely serious adverse effects by matching those targets to proteins implicated in specific side-effect phenotypes. To achieve this, we developed a boosted tree regression machine learning approach that uses the above 6,400 proteins as a base feature set. If a drug binds to a protein, the corresponding vector component value is set to the frequency of that protein multiplied by a factor based on binding mode (see Methods); otherwise, it is set to zero. For 1,079 DrugBank drugs that mapped to the SIDER4 dataset^2^, from which drug’s serious side effects were obtained, the AUC is 0.68. With a cutoff score of 0.39 that maximizes the Matthews correlation coefficient (MCC=0.29), the recall is 54% and the precision is 52%. For the same set of drugs and serious side-effects at the maximal MCC=0.24, the recall of MEDICASCY^7^ is 39% and the precision is 53%. Thus, for roughly the same precision, PHENOCAUZ achieves ∼15% more recall.

Together, these analyses show how a molecular symptom map can be converted into an interpretable target-safety framework. PHENOCAUZ does not infer toxicity from chemical structure alone; instead, it uses target identity, binding mode and the phenotype associations of those targets to propose mechanistic links between pharmacological perturbation and serious adverse outcomes. This target-centered representation is complementary to structure-based predictors such as MEDICASCY^7^.

### Identification of candidate therapeutics to alleviate adverse symptoms

From DrugBank^24^, we obtained 17,430 drugs and their recorded protein interactions. Using both OMIM-derived and predicted candidate symptom-driver proteins, we screened all 7,152 symptoms against these drugs across all recorded binding modes. For 3,134 drugs, exact name matching mapped indications to 1,356 symptoms, including 30 cancers. Across these 1,356 symptoms, the screen yielded a mean enrichment factor of 5.2 and mean recall of 45%. These are drug-retrieval metrics, not estimates of recovery through established therapeutic targets: the interaction set was not restricted to pharmacodynamic targets and can include proteins involved in drug absorption, distribution, metabolism and excretion (ADME). The predicted drug lists can be found in Supplementary Materials, Table S9.

### Examples

In what follows, we examine in more detail the predicted candidate treatments for ovarian, prostate, and breast cancers, dementia and Crohn’s disease.

### Application to cancer drug discovery

#### Ovarian cancer

PHENOCAUZ identified 1,838 putative ovarian cancer-associated proteins (324 OMIM-derived and 1,514 predicted; see Supplementary Materials, Tables S2 and S6), of which 241 overlap the 723 COSMIC census genes^36^. Screening these proteins against 17,430 DrugBank drugs yielded 2,982 candidates (see Supplementary Materials, Table S9). The screen retrieved 16 of 43 drugs annotated for ovarian cancer (37% recall; enrichment factor 2.2). Retrieved drugs included paclitaxel, which stabilizes microtubules, and doxorubicin, which intercalates DNA and poisons topoisomerase II, including TOP2A. The screen also linked vinorelbine, trabectedin and rucaparib to CYP2D6. Such links should not be interpreted as validation of CYP2D6 as an ovarian cancer driver or therapeutic target but suggests it might be a novel target.

#### Prostate cancer

PHENOCAUZ identified 2,072 prostate cancer-associated proteins (235 OMIM-derived and 1,837 predicted), including 215 of the 723 COSMIC census genes^36^. The screen returned 2,869 candidates (see Supplementary Materials, Table S9) and retrieved 36 of 86 drugs annotated for prostate cancer giving 42% recall and an enrichment factor 2.5. Of the 36 retrieved drugs, 21 had recorded interactions with OMIM-derived proteins and 15 with predicted proteins. For example, Ridaforolimus provides an example of mTOR inhibition^37^. Another drug is Rucaparib which is a PARP inhibitor^38^. Lutetium Lu-177 vipivotide tetraxetan, a PSMA-targeted radioligand, was also retrieved^39^.

#### Breast cancer

PHENOCAUZ identified 2,302 breast cancer-associated proteins, including 267 of the 723 COSMIC census genes. The screen yielded 3,964 drugs (see Supplementary Materials, Table S9), an enrichment factor of 3.0 and recall of 68% (75/111). Of the 75 retrieved drugs, 58 had recorded interactions with OMIM-derived proteins and 17 with predicted proteins. Examples include the CDK4/6 inhibitor ribociclib and the estrogen-receptor antagonist fulvestrant^40^. Vinorelbine was also retrieved^41^, but its antitumor mechanism is disruption of microtubule assembly. Lasofoxifene was another retrieved drug^42^. As in the ovarian and prostate cancer analyses, the reported metrics describe the interaction screen rather than recovery exclusively through established therapeutic targets.

#### Application to Dementia

PHENOCAUZ identified 3,633 putative dementia-associated proteins (784 OMIM-derived and 2,849 predicted). The unfiltered DrugBank screen yielded 4,771 candidates (see Supplementary Materials, Table S9) and retrieved 14 of 22 drugs annotated for dementia (64% recall; enrichment factor 2.3). Since testing all the drugs for efficacy is beyond this work, the majority of the retrieved drugs have unknown for efficacy for Dementia treatment. Five retrieved drugs had recorded interactions with OMIM-derived proteins and nine with predicted proteins. Retrieved examples included lecanemab^43^, memantine^44^, nicergoline^45^ and galantamine. Again, these retrieval results do not by themselves validate the individual protein links or establish efficacy across dementia subtypes; therapeutic mechanisms and indication-specific evidence require separate experimental assessment.

#### Application to Crohn’s disease

We aggregated 384 (194 known, 190 novel) driver proteins for Crohn’s disease (CD). This yielded 1,731 putative efficacious drugs after screening against DrugBank drugs (see Supplementary Materials, Table S9). Of these, 13 of the 24 known efficacious drugs for Crohn’s disease listed in DrugBank were recalled (54%) with an enrichment factor of 5.5. Examples of drugs binding to known targets include: fontolizumab binding IFNG, which has been investigated for treatment in CD^46^; another metronidazole, a CYP2C8 inhibitor, has been shown to be efficacious for specific manifestations and complications of Crohn’s disease^47^. Examples of drugs binding to predicted targets include briakinumab, an IL12B binder, that has been investigated for treating CD^48^; PN0621 binding to TNF, is an experimental drug intended for treating CD^24^.

#### Summary of PHENOCAUZ based drug-disease prediction results

The inferred/predicted symptom–protein relationships were used to identify potential therapeutic agents. Screening symptom-causing proteins against DrugBank drugs yielded candidate treatments for ovarian, prostate, and breast cancers with recall rates of 37%, 42%, and 68%, respectively. Additional predictions identified potential therapeutics for dementia and Crohn’s disease with recall rates of 64% and 54%, respectively. Recalled drugs include paclitaxel for ovarian cancer; rucaparib for prostate cancer; ribociclib for breast cancer; lecanemab for dementia; and briakinumab for Crohn’s disease.

#### Application to multiple-target-based drug discovery

The above examples show how one can target the causative proteins themselves to discover efficacious drugs, especially from the approved and experimental drugs in DrugBank that have experimentally validated targets and clear molecular mechanisms. These drugs can be repurposed for new indications. We note that in cases where the causative proteins are not expressed, or are undruggable, alternative approaches are sought. In earlier work^49^, we developed a multiple-target based method called CoVLS using predicted drug targets for new drug discovery as well as drug repurposing. The CoVLS approach identifies efficacious drugs by FINDSITE^comb2.0 27^ virtual ligand screening of the comorbidity frequency weighted input MOA proteins for a given adverse response. A drug is ranked by the dot product of the predicted molecule’s binding precision and the p-value weighted frequency that the protein is a MOA protein of a comorbid disease, normalized by the summed binding precision of all its human targets^49^. Here we test a scenario that uses PHENOCAUZ causative proteins’ pathways to filter the LeMeDISCO-derived mode-of-action (MOA) proteins (i.e., the MOAs must be in the same pathways as those involved by causative proteins) and screens against the subset of 2,095 FDA-approved DrugBank drugs, all of which we have predicted targets. On a set of 346 symptoms mapped to LeMeDISCO’s diseases and having non-zero efficacious FDA-approved drugs, the mean recall rate, observed precision (correct prediction/total prediction) and enrichment factor are 50.5%, 0.87% and 3.1, respectively, compared to 51.1%, 0.80% and 2.7 without the pathway filter. Although the improvement by PHENOCAUZ’s pathway protein filtering is small, it shows that by ignoring MOA proteins not in the same pathways as the causative proteins, the prediction performance does not deteriorate. We note that the mean recall rate of 50.5% by pathway filtered CoVLS and 51.1% by pure CoVLS are even slightly better than the 45% recall obtained from the above screening based on one or a few experimental targets. We also tested a scenario that uses all the proteins involved in the PHENOCAUZ causative proteins’ pathways, the recall rate, observed precision, and enrichment factor were 81.9%, 0.3%, and 1.08, respectively. Although the recall rate is higher; the observed precision and enrichment factor (slightly better than random) are much lower than those of the LeMeDISCO’s MOAs and MOAs filtered by PHENOCAUZ protein pathways. This indicates that not all PHENOCAUZ protein pathways’ proteins are efficacious targets and only a subset that overlap with LeMeDISCO’s MOAs are efficacious. Combining LeMeDISCO’s MOAs and PHENOCAUZ’s protein pathway-filter is a more effective approach than either pure LeMeDISCO’s MOAs or PHENOCAUZ’s pathway proteins for multiple-target-based drug discovery. Future work will tease out the molecular features that distinguish efficacious and non-efficacious targets in the same pathway.

Finally, we note that a mean enrichment factor of 8.5 of drugs was found to treat 7,152 symptoms; indeed, 82.7% of symptoms had enrichment factors greater than 1. For example, for anxiety disorder, 16 of the 17 known drugs that treat this disease were recalled from 743 drug predictions with an enrichment factor (EF) of 2.7; 42 of 46 asthma drugs were recalled from 890 drug predictions with an EF of 2.1. For liver cirrhosis, the 37 predicted drugs recalled 5 of 8 known drugs with an EF of 35. Thus, PHENOCAUZ filtering provides better precision and enrichment factor while causing only a very small decrease in the recall rate (relatively ∼1%).

## Discussion

PHENOCAUZ treats Mendelian disorders as natural human perturbation experiments from which the molecular architecture of clinical phenotypes can be learned. By integrating Mendelian genotype-phenotype relationships with pathway, biological-process and disease-propensity features, the framework maps thousands of symptoms to molecular contexts and extends those relationships to candidate symptom drivers across the human proteome. Independent curated resources support many of these predictions directly and show even stronger agreement at the pathway level. These results support the broader premise that clinical phenotypes contain recoverable information about the molecular systems whose perturbation produces them.

A central result is the distinction between proteins implicated in generating a symptom and proteins through which that symptom can be therapeutically modified. Efficacious drug targets and LeMeDISCO mode-of-action proteins show only modest direct overlap with PHENOCAUZ symptom drivers, yet their pathways overlap for more than 90% of evaluated symptoms. Therefore, an effective therapy need not act directly on a primary disease driver; it can act through upstream regulators, downstream effectors or compensatory nodes within the same molecular system. This pathway-level convergence provides a mechanistic explanation for why therapeutic targets can be biologically close to a causal lesion without being identical to it.

The same phenotype-to-mechanism representation could have practical applications in biomedical engineering and drug development. Proteins linked to severe phenotypes define interpretable candidate risk targets, and target identity plus binding mode improves recall of serious adverse drug reactions relative to our earlier structure-based approach at comparable precision. Conversely, mapping disease-associated symptom drivers to experimentally characterized drug targets prioritizes candidate therapeutics across cancer, dementia and Crohn’s disease. These applications should be viewed as demonstrations of the framework rather than proof that every predicted protein is causal or every nominated drug is efficacious; prospective experimental and clinical validation will be required for individual predictions. More broadly, PHENOCAUZ provides a scalable route from clinical phenotype to molecular mechanism and from mechanism to testable therapeutic and safety hypotheses. CoVLS screening tests indicate that a subset of proteins in the pathways of PHENOCAUZ drivers are efficacious targets. Thus, this approach should be viewed as symptom-protein-drug target prioritization approach that could hold the promise of accelerating drug discovery.

## Acknowledgments

This research was supported in part by NIH grant GM-118039 by gift from the Ovarian Cancer Institute. We thank Bartosz Ilkowski for computational support Jessica Forness for assistance with manuscript preparation and proofreading.

## Data and Code Availability

Scripts and data required to reproduce the prediction results are available at: https://github.com/hzhou3ga/symptomprotein/

## Author contributions

Conceptualization: JS, HZ

Data curation: HZ

Formal analysis: HZ

Funding acquisition: JS

Investigation: HZ, JS

Methodology: HZ

Project administration: JS

Resources: JS

Software: HZ

Supervision: JS

Validation: HZ, JS

Writing – original draft: HZ

Writing – review & editing: JS, HZ

## Materials and Methods

An overview of PHENOCAUZ is presented in Figure 1 with the detailed steps explained below.

### Inferring protein-symptom causative relationships

The OMIM database^3^ provides Mendelian protein-phenotype causative relationships. For each phenotype, we used the PubMed E-utilities^11^ API to search for publications with keywords from both phenotypes and symptoms independently. A total of 8,722 phenotypes were searched. The symptoms were collected from the 16,190 symptom terms in SympGAN^5^, plus the set of 4,251 side effect terms from the SIDER (version 4) database^2^. Note that 621 terms in these two symptom sets overlap. After removing rare symptoms whose number keywords > 10, we obtained 15,710 non-redundant symptom terms. E-utilities return a list of publications containing the set of keywords. Word order was unrestricted, but the terms were required to occur next to each other within title or abstract fields, specifically with syntax “[tiab:∼0]”. Subsequently, for each phenotype-symptom pair, the strength of association was measured using the Jaccard index (J-score)^50^ and a p-value computed using Fisher’s exact test^51^. For a given phenotype, the symptom p-values are then converted to q-values to control the false discovery rates. With a cutoff of q-value < 0.05, 7,152 symptoms are present in at least one phenotype having corresponding causative gene annotation. The resulting list of symptoms are provided in Supplementary Materials, Table S1. Their phenotypes are then mapped to 4,222 non-redundant proteins using the OMIM^3^ dataset to obtain protein-symptom relationships (Supplementary Materials, Table S2 provides the complete list without predicted proteins). The resulting protein-symptom list will be applied in AI model training and in identifying important pathways responsible for a given symptom.

### Predicting protein-symptom causative relationships

The above inferred protein-symptom relationships cover only those proteins associated with Mendelian phenotypes. To account for the possible contribution of proteins not known to be associated with a Mendelian disease, we developed an AI model trained on these known causative protein-symptom data by learning the protein features that contribute most to a specific symptom. We next describe how PHENOCAUZ accomplishes this goal.

### Calculation of protein disease-causing propensity

To account for the protein’s disease/phenotype-causing propensity, we include the protein’s aggregate deleterious variation score from the average pathogenicity score (i.e. the summed score normalized by the total number of possible variations) of all possible exome variations of a protein obtained from our state-of-the-art ENTPRISE^52^ and ENTPRISE-X^53^ algorithms that predict disease-associated variants. For ENTPRISE’s evaluation of missense variations, all wildtype exome residue positions are mutated to all other 19 amino acid types. Then, their variation scores are averaged to provide the aggregate score for a given protein. Similarly, in ENTPRISE-X’s evaluation of nonsense variations, all residue positions are assumed to be mutated to a stop codon, and the resulting scores averaged.

### Assignment of a protein to its pathways and GO processes

Pathways and their corresponding associated proteins are obtained from the Reactome database^12^ which contains 2,363 distinct pathways. If a protein is present in a pathway, the protein component corresponding to this pathway is set to 1; otherwise, it is set to 0. The total number of pathways in which a given protein is involved is also included as a feature component.

The 12,535 unique human biological processes represented in GO annotations were downloaded from http://proteinontology.org/docs/download-ontology/^18^. If a protein is involved in a GO process, the component corresponding to that process is set to 1, otherwise, it is set to 0. The total number of GO processes in which a given protein is involved is also included.

Concatenating all the above features leads to a 14,902-dimensional feature vector for each of the 18,409 unique proteins present in Reactome pathways and GO processes.

### Machine learning

We employed the Boosted Random Forest (BRF) regression machine learning method developed in our earlier work, MEDICASCY^7^. A BRF is an iterated application of Random Forests in which, at each iteration, the residual function value from all previous fits, *RF_n_*(*X*) is fitted:

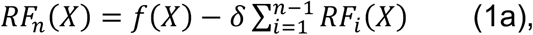

where *δ* is a learning rate parameter. In this work, we empirically set *δ* = 0.2 and the total number of iterations to be N_RF_=25. The final regression function is

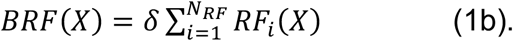

We implement the BRF using the standard Python package scikit-learn (https://scikit-learn.org/) with the default setting (*n_estimators*=10) of the multi-value regression function RandomForestRegressor. For training, the multilabel objective values of each gene are their J-scores computed for the associations between its phenotype and the 7,152 symptoms. For all predictions, we convert the raw score *S*_0_ from machine learning to the normalized precision score (ranging from 0 to 1) by:

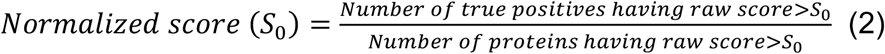

### Prediction of serious side effects from drug targets

Drug targets and binding modes were obtained from DrugBank v5.0^24^ and their corresponding serious side effects were taken from the SIDER4 dataset^2^. For a given drug, the feature vector contains 6,400 components for the list of proteins causing serious symptoms. If a protein is not targeted by the drug, the component value is defaulted to 0; if the drug binds to a protein as activator, inducer, agonist, stimulator, chaperone, or upregulator, the component value is also set to 0 based on preliminary testing, which indicated that 0 outperformed other non-zero values. However, if the binding mode is one of the following function-reducing modes: antagonist, inhibitor, neutralizer, suppressor, blocker, antibody, downregulator, inactivator, chelator, degradation, adduct, the component value is set to be the *negative* value of the serious symptom frequency of the protein; for proteins with any other binding modes, the component values are set to the frequencies of the proteins. Two additional feature components represent the total numbers of function-reducing targets inside and outside the 6,400 protein list that cause serious symptoms. As in the ENTPRISE work^52,54^, a boosted tree regression machine learning method was employed for training and prediction. Here, a decision tree regression is implemented with a maximum depth of 6. The scoring function is represented as a boosted decision tree:

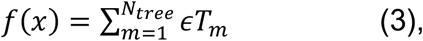

where *T_m_* is a decision tree, *ε* is the shrinkage factor or learning rate, and *N_tree_* is the number of trees. The learning rate *ε* is set to 0.05 and the number of trees, *N_tree_* is set to 2000. These values are purely empirical and not optimized.

## References

1 Alghamdi, J. & Padmanabhan, S. in Handbook of Pharmacogenomics and Stratified Medicine (ed Sandosh Padmanabhan) 235–257 (Academic Press, 2014).

2 Kuhn, M., Letunic, I., Jensen, L. J. & Bork, P. The SIDER database of drugs and side effects. Nucleic Acids Res 44, D1075–1079 (2016). 10.1093/nar/gkv1075

3 McKusick, V. A. Online Mendelian Inheritance in Man, OMIM®.

4 Zhou, X., Menche, J., Barabási, A.-L. & Sharma, A. Human symptoms– disease network. Nature Communications 5, 4212 (2014).

5 Lu, K. et al. SympGAN: A systematic knowledge integration system for symptom–gene associations network. Knowledge-Based Systems 276, 110752 (2023). 10.1016/j.knosys.2023.110752

6 Miaskowski, C. et al. Advancing Symptom Science Through Symptom Cluster Research: Expert Panel Proceedings and Recommendations. JNCI: Journal of the National Cancer Institute 109, djw253 (2017). 10.1093/jnci/djw253

7 Zhou, H. et al. MEDICASCY: A Machine Learning Approach for Predicting Small-Molecule Drug Side Effects, Indications, Efficacy, and Modes of Action. Molecular Pharmaceutics 17, 1558–1574 (2020). 10.1021/acs.molpharmaceut.9b01248

8 Muñoz, E., Novácek, V. & Vandenbussche, P. Y. Facilitating prediction of adverse drug reactions by using knowledge graphs and multi-label learning models. Brief Bioinform 20, 190–202 (2019). 10.1093/bib/bbx099

9 Dimitri, G. M. & Lió, P. DrugClust: A machine learning approach for drugs side effects prediction. Comput Biol Chem 68, 204–210 (2017). 10.1016/j.compbiolchem.2017.03.008

10 Zhong, Y., Seoighe, C. & Yang, H. Non-Negative matrix factorization combined with kernel regression for the prediction of adverse drug reaction profiles. Bioinform Adv 4, vbae009 (2024). 10.1093/bioadv/vbae009

11 Kans, J. E-utilities on the Unix Command Line. Entrez® Programming Utilities Help [Internet] 2010-. (2013).

12 Jassal, B. et al. The reactome pathway knowledgebase. Nucleic Acids Res 48, D498–d503 (2020). 10.1093/nar/gkz1031

13 Cao, B. et al. Pathology of pain and its implications for therapeutic interventions. Signal Transduction and Targeted Therapy 9, 155 (2024). 10.1038/s41392-024-01845-w

14 McEntire, D. M. et al. Pain transduction: a pharmacologic perspective. Expert Rev Clin Pharmacol 9, 1069–1080 (2016). 10.1080/17512433.2016.1183481

15 Duman, R. S. & Voleti, B. Signaling pathways underlying the pathophysiology and treatment of depression: novel mechanisms for rapid-acting agents. Trends Neurosci 35, 47–56 (2012). 10.1016/j.tins.2011.11.004

16 Wen, X. et al. Signaling pathways in obesity: mechanisms and therapeutic interventions. Signal Transduction and Targeted Therapy 7, 298 (2022). 10.1038/s41392-022-01149-x

17 Zhang, Y. R. et al. Immune-mediated diseases are associated with a higher incidence of dementia: a prospective cohort study of 375,894 individuals. Alzheimers Res Ther 14, 130 (2022). 10.1186/s13195-022-01072-x

18 Ashburner, M. et al. Gene Ontology: tool for the unification of biology. Nature Genetics 25, 25–29 (2000). 10.1038/75556

19 Fan, Y. & Flannick, J. A Novel Algorithm for Rare Disease Gene Prediction Based on Phenotypic Similarity. Journal of the Endocrine Society 5, A497– A498 (2021). 10.1210/jendso/bvab048.1017

20 Zheng, W. et al. PERADIGM: Phenotype embedding similarity-based rare disease gene mapping. PLoS Genet 21, e1011976 (2025). 10.1371/journal.pgen.1011976

21 Falaguera, M. J. et al. Temporal trends in evidence supporting novel drug target discovery. Nat Commun 17, 492 (2026). 10.1038/s41467-025-67180-y

22 Bodenreider, O. The Unified Medical Language System (UMLS): integrating biomedical terminology. Nucleic Acids Res 32, D267–270 (2004). 10.1093/nar/gkh061

23 Schriml, L. M. et al. Disease Ontology: a backbone for disease semantic integration. Nucleic Acids Research 40, D940–D946 (2012). 10.1093/nar/gkr972

24 Wishart, D. S. et al. DrugBank 5.0: a major update to the DrugBank database for 2018. Nucleic Acids Res 46, D1074–D1082 (2018). 10.1093/nar/gkx1037

25 Zhang, Y. et al. Therapeutic target database 2026: facilitating targeted therapies and precision medicine. Nucleic Acids Res 54, D1692–d1701 (2026). 10.1093/nar/gkaf1154

26 Astore, C., Zhou, H., Ilkowski, B., Forness, J. & Skolnick, J. LeMeDISCO is a computational method for large-scale prediction & molecular interpretation of disease comorbidity. Communications Biology 5, 870 (2022).

27 Zhou, H., Cao, H. & Skolnick, J. FINDSITE^comb2.0^: A New Approach for Virtual Ligand Screening of Proteins and Virtual Target Screening of Biomolecules. Journal of Chemical Information and Modeling 58, 2343– 2354 (2018).

28 Valderas, J. M., Starfield, B., Sibbald, B., Salisbury, C. & Roland, M. Defining comorbidity: implications for understanding health and health services. Annals of family medicine 7, 357–363 (2009).

29 Jenewein, T. et al. Genotype-phenotype dilemma in a case of sudden cardiac death with the E1053K mutation and a deletion in the SCN5A gene. Forensic Sci Int 275, 187–194 (2017). 10.1016/j.forsciint.2017.02.038

30 Einspahr, J. M. et al. Cardiomyocyte-specific deletion of adhesion G protein-coupled receptor F5 (ADGRF5) increases susceptibility to stress and arrhythmia. The Journal of Pharmacology and Experimental Therapeutics 389, 64 (2024). 10.1124/jpet.064.941530

31 Zhao, R., Choi, B. Y., Lee, M. H., Bode, A. M. & Dong, Z. Implications of Genetic and Epigenetic Alterations of CDKN2A (p16(INK4a)) in Cancer. EBioMedicine 8, 30–39 (2016). 10.1016/j.ebiom.2016.04.017

32 Cui, Z., Kang, H., Grandis, J. R. & Johnson, D. E. CYLD Alterations in the Tumorigenesis and Progression of Human Papillomavirus-Associated Head and Neck Cancers. Mol Cancer Res 19, 14–24 (2021). 10.1158/1541-7786.Mcr-20-0565

33 Mungrue, I. N. et al. Abcc6 deficiency causes increased infarct size and apoptosis in a mouse cardiac ischemia-reperfusion model. Arterioscler Thromb Vasc Biol 31, 2806–2812 (2011). 10.1161/atvbaha.111.237420

34 Śniadecki, M0et al. BARD1 and Breast Cancer: The Possibility of Creating Screening Tests and New Preventive and Therapeutic Pathways for Predisposed Women. Genes (Basel) 11 (2020). 10.3390/genes11111251

35 Gao, Y., Chen, F., Niu, S., Lin, S. & Li, S. Replication and Meta-Analysis of Common Gene Mutations in TTF1 and TTF2 with Papillary Thyroid Cancer. Medicine (Baltimore) 94, e1246 (2015). 10.1097/md.0000000000001246

36 Tate, J. G. et al. COSMIC: the Catalogue Of Somatic Mutations In Cancer. Nucleic Acids Res 47, D941–d947 (2019). 10.1093/nar/gky1015

37 Squillace, R. M. et al. Synergistic activity of the mTOR inhibitor ridaforolimus and the antiandrogen bicalutamide in prostate cancer models. Int J Oncol 41, 425–432 (2012). 10.3892/ijo.2012.1487

38 Bryce, A. H., Nash, P., Roessler, A. & Mandair, D. Rucaparib - drug evaluation: a PARP inhibitor for the treatment of BRCA-mutated metastatic castration-resistant prostate cancer. Future Oncol 22, 1759–1773 (2026). 10.1080/14796694.2026.2681674

39 Zhang, A. Y., Bertsch, H., Chaudhary, A. & Salner, A. Lutetium (177)Lu Vipivotide Tetraxetan Efficacy and Toxicity in Advanced Prostate Cancer. Adv Radiat Oncol 11, 101917 (2026). 10.1016/j.adro.2025.101917

40 McKeage, K., Curran, M. P. & Plosker, G. L. Fulvestrant: a review of its use in hormone receptor-positive metastatic breast cancer in postmenopausal women with disease progression following antiestrogen therapy. Drugs 64, 633–648 (2004). 10.2165/00003495-200464060-00009

41 Cybulska-Stopa, B. et al. Evaluation of vinorelbine-based chemotherapy as the second or further-line treatment in patients with metastatic breast cancer. Contemp Oncol (Pozn) 17, 78–82 (2013). 10.5114/wo.2013.33779

42 Gal-Yam, E. N. & Levanon, K. Lasofoxifene Monotherapy Induces Durable Complete Remission in a Patient with Estrogen Receptor-Positive, Metastatic Breast Cancer with an ESR1 Mutation. JCO Precision Oncology, e2300097 (2023). 10.1200/po.23.00097

43 Honig, L. S. et al. Updated safety results from phase 3 lecanemab study in early Alzheimer’s disease. Alzheimers Res Ther 16, 105 (2024). 10.1186/s13195-024-01441-8

44 McShane, R. et al. Memantine for dementia. Cochrane Database Syst Rev 3, Cd003154 (2019). 10.1002/14651858.CD003154.pub6

45 Fioravanti, M. & Flicker, L. Efficacy of nicergoline in dementia and other age associated forms of cognitive impairment. Cochrane Database Syst Rev 2001, Cd003159 (2001). 10.1002/14651858.Cd003159

46 Reinisch, W. et al. Fontolizumab in moderate to severe Crohn’s disease: a phase 2, randomized, double-blind, placebo-controlled, multiple-dose study. Inflamm Bowel Dis 16, 233–242 (2010). 10.1002/ibd.21038

47 Babb, R. R. The use of metronidazole (Flagyl) in Crohn’s disease. J Clin Gastroenterol 10, 479–481 (1988). 10.1097/00004836-198810000-00002

48 Panaccione, R. et al. Briakinumab for treatment of Crohn’s disease: results of a randomized trial. Inflamm Bowel Dis 21, 1329–1340 (2015). 10.1097/mib.0000000000000366

49 Astore, C., Zhou, H., Jacob, J. & Skolnick, J. Prediction of severe adverse events, modes of action and drug treatments for COVID-19’s complications. Scientific Reports 11, 20864 (2021). 10.1038/s41598-021-00368-6

50 THE DISTRIBUTION OF THE FLORA IN THE ALPINE ZONE. (1912).

51 Fisher, R. A. On the Interpretation of χ 2 from Contingency Tables, and the Calculation of P. Journal of the Royal Statistical Society 85, 87–94 (1922).

52 Zhou, H., Gao, M. & Skolnick, J. ENTPRISE: An Algorithm for Predicting Human Disease-Associated Amino Acid Substitutions from Sequence Entropy and Predicted Protein Structures. PLoS One 11, e0150965 (2016). 10.1371/journal.pone.0150965

53 Zhou, H., Gao, M. & Skolnick, J. ENTPRISE-X: Predicting disease-associated frameshift and nonsense mutations. PLoS One 13, e0196849 (2018). 10.1371/journal.pone.0196849

54 Friedman, J. H. & Meulman, J. J. Multiple additive regression trees with application in epidemiology. Statistics in Medicine 22, 1365–1381 (2003). 10.1002/sim.1501

